# Human *METTL7B* Encodes an Alkyl Thiol Methyltransferase that Methylates Hydrogen Sulfide

**DOI:** 10.1101/2020.03.06.979542

**Authors:** Benjamin J. Maldonato, Rheem A. Totah

**Author notes:** Corresponding Author: Dr. Rheem A. Totah.

## Abstract

Methyltransferase-like protein 7B (METTL7B) is implicated in tumor growth and progression while gene expression is upregulated in several different disease states such as rheumatoid arthritis and breast cancer. Yet, the catalytic function of METTL7B has not been characterized. Here we demonstrate that *METTL7B* encodes a protein that catalyzes the transfer of a methyl group from S-adenosyl-L-methionine (SAM) to hydrogen sulfide (H_2_S) to form methanethiol (CH_3_SH). Several exogenous aliphatic thiols were also identified as substrates. Modulation of *METTL7B* gene expression in HepG2 and HeLa cell culture directly alters the methylation of captopril, a marker reaction of alkyl thiol methyltransferase (TMT) activity(1, 2). Furthermore, cloned and recombinantly expressed and purified METTL7B full length protein methylates several thiol compounds, including hydrogen sulfide, 7α-thiospironolactone, captopril, and L-penicillamine in a concentration dependent manner. Endogenous thiols such as glutathione and cysteine or classic probe substrates of other known small molecule *S*-, *N*-, and *O*- methyltransferases were not substrates for METTL7B. Our results unequivocally demonstrate, and for the first time, that METTL7B, a protein implicated in several disease states, is an alkyl thiol methyltransferase(3–5). Identifying the catalytic function of METTL7B will enable future pharmacological research in disease pathophysiology where *METTL7B* expression and H_2_S levels can potentially alter the redox state and growth cycle of cells.

## Introduction

Hydrogen sulfide (H_2_S) is a gasotransmitter that regulates inflammatory and cell cycle processes(6). It is biosynthesized by three different enzymes, cystathionine γ-lyase (CSE), cystathionine ß-synthase (CBS), and 3-mercaptopyruvate sulfurtransferase (3-MST)(7). CBS activity is inhibited by carbon monoxide and nitric oxide but is activated by S-adenosyl-L-methionine (SAM)(8, 9). Therefore, production of hydrogen sulfide is sensitive to intracellular redox state. Once formed, hydrogen sulfide causes physiological effects by formation of persulfide bonds to protein cysteine residues(10). Catabolism of hydrogen sulfide is believed to be primarily driven by oxidation(11). This route of metabolism may be less prominent in organs outside of the gut and under hypoxic conditions, such as in the interior of solid tumors(12, 13). In these instances, methylation can play a key role in hydrogen sulfide catabolism yet little is known about this process or the enzyme that catalyzes this reaction. In this report, we identify METTL7B as an alkyl thiol methyltransferase that catalyzes the transfer of a methyl group from S-adenosyl-L-methionine (SAM) to hydrogen sulfide and several aliphatic thiol-containing compounds.

To date, the catalytic function of methyltransferase-like protein 7B (METTL7B) was unknown despite being implicated in several disease states. Specifically, *METTL7B* gene expression is significantly altered in kidney disease, acute respiratory distress syndrome, and numerous cancers, including breast, non-small cell lung, thyroid, and ovarian(3–5, 14–17). METTL7B expression appears to be responsive to inflammation signaling pathways via JAK1(18, 19). Gene expression also changes with respect to cellular redox state and is associated with individual response to certain chemotherapeutics(20, 21). In non-small cell lung cancer, *METTL7B* contributes to tumorigenesis and progression by regulating cell cycle progression. Gene silencing reduced tumor growth and progression both *in vitro* and *in vivo* suggesting METTL7B as a potential therapeutic target(17).

Interest in METTL7B originated as we were attempting to identify the elusive alkyl thiol methyltransferase (TMT) responsible for the methylation of the active metabolite of clopidogrel *in vivo*(22). This microsomal enzyme catalyzes the methylation of aliphatic thiols in humans, including hydrogen sulfide, captopril, 7α-thiospironolactone, D- and L-penicillamine, and the active metabolites of prasugrel, and ziprasidone(1, 2, 23–27). However, despite numerous attempts, researchers have not successfully identified the TMT gene or protein (28–31).

Our preliminary approach to identify TMT expanded on earlier research which attempted to purify TMT from rat liver microsomes using a number of chromatographic steps(28, 31). After significant increases in TMT specific activity, preliminary non-targeted proteomic experiments were conducted to identify potential methyltransferase proteins in the TMT-active fractions. The major candidate protein in active fractions was identified as rat METTL7B which was also localized to the endoplasmic reticulum (Extended Data Table 1). Rat and human METTL7B share 83% sequence homology, which suggests a conserved function. Subsequent experiments modulating the expression of human METTL7B in two cell lines also altered captopril methylation, a known TMT substrate. Once identified, we cloned, recombinantly expressed, and purified human full length METTL7B in *E. coli* and conducted small molecule substrate screening with the purified protein. The activity screens confirmed that METTL7B specifically catalyzes SAM-dependent methylation of aliphatic thiol compounds, including hydrogen sulfide, in a time and protein concentration dependent manner. No methylation was observed with classic probe substrates of other known small molecule *S*-, *N*-, and *O*- methyltransferases(32–36) or endogenous thiols such as cysteine or glutathione.

## Results

### METTL7B Gene Expression Modulation in Mammalian Cell Culture

Treating HepG2 cells with *METTL7B* specific small interfering RNA (siRNA) caused an average of 60% decrease in *METTL7B* mRNA expression compared to cells treated with a scrambled negative siRNA control (Figure 1A). Incubation with captopril following siRNA treatment, a previously reported TMT- probe substrate, showed an average 51% decrease in captopril methylation in HepG2 cells with reduced *METTL7B* gene expression (Figure 1B).

**Figure 1.**
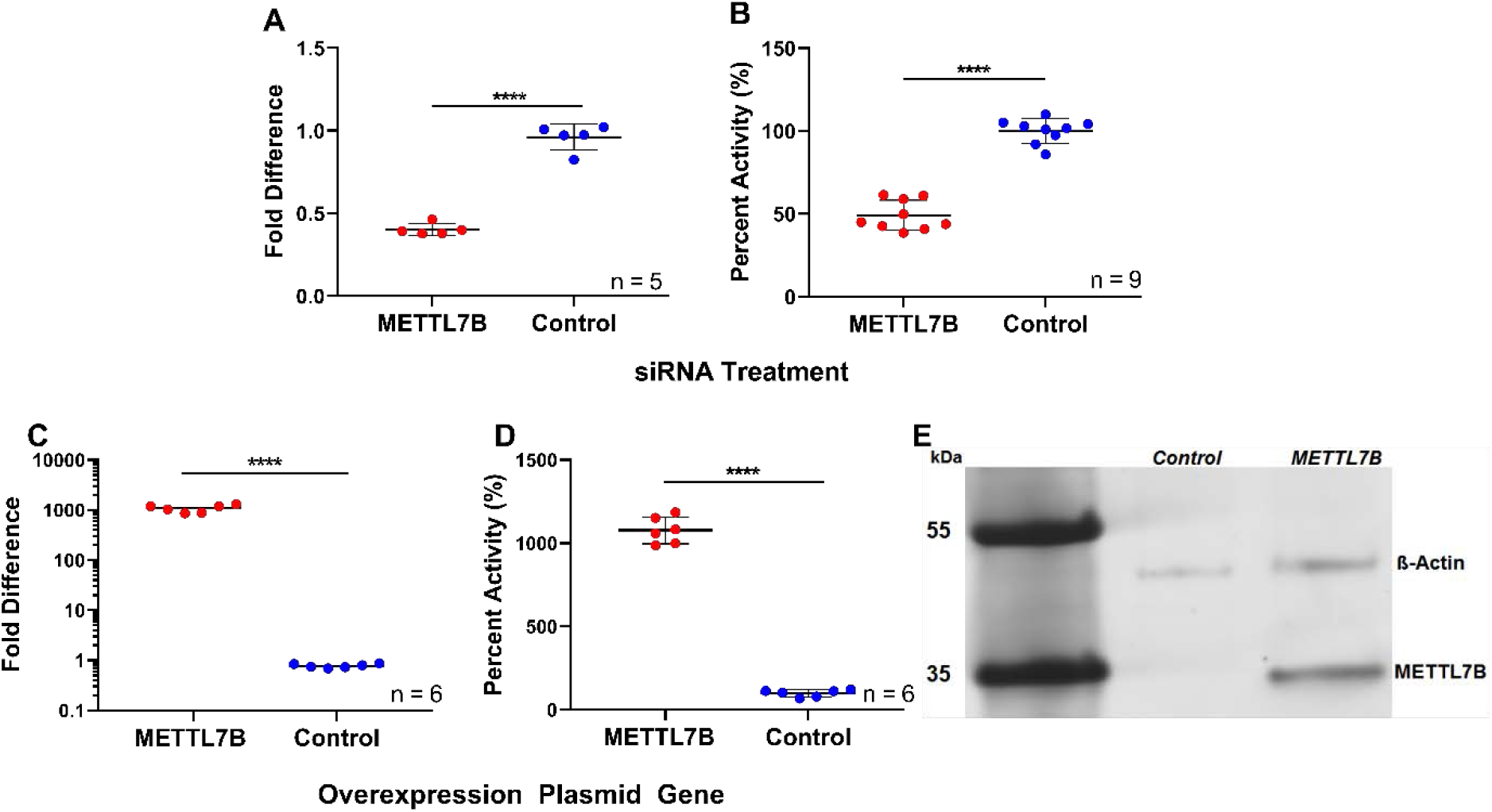
Alteration of *METTL7B* gene expression in human cell culture: **A)** *METTL7B* gene expression significantly decreased in HepG2 cells compared to controls when treated with anti-*METTL7B* siRNA for 72 hours. **B)** Methylation of captopril activity significantly decreased in HepG2 cells treated with *METTL7B* siRNA compared to control cells. **C)** HeLa cells treated with a *METTL7B* overexpression plasmid showed ~ 1,000-fold increase in *METT7B* gene expression compared to control cells transfected with an empty expression vector. **D)** HeLa cells transfected with the *METTL7B* expression vector showed a 10-fold increase in captopril methylation activity compared to negative control cells. **E)** FLAG-tagged METTL7B expression is only observed in cells treated with the *METTL7B* overexpression plasmid compared to controls (Lanes 2 and 1 respectively). ß-actin was used as a loading control. All data is presented as the mean ± standard deviation. Individual data points from two (A, C, and D) or three (B) experiments are plotted. Significance was determined using unpaired two-tailed *t* test. \*\*\*\**P*<0.0001.

*METTL7B* gene expression increased over 1,000-fold in HeLa cells treated with a constitutive overexpression plasmid containing the *METTL7B* gene sequence compared to cells treated with an empty control plasmid as measured by RT-PCR (Figure 1C). Captopril methylation subsequently increased 10-fold in cells overexpressing *METTL7B* compared to control cells (Figure 1D). Cells transfected with the METTL7B overexpression plasmid show formation of FLAG-tagged METTL7B (Lane 2, Figure 1E) compared to cells treated with an empty overexpression plasmid (Lane 1, Figure 1F).

### Expression and Purification of METTL7B Fusion Protein

The full-length *METTL7B* gene sequence was inserted into a pET21 expression plasmid to express a unique fusion protein in *E. coli*. The fusion protein, henceforth referred to as pET21 METTL7B, is 57.5 kDa and contains a dual His-GST affinity/solubilization tag coupled to the N-terminus of the native METTL7B protein. We developed a dual-stage affinity purification protocol as detailed in the Methods section. The resulting purified protein fraction predominantly contains the pET21 METTL7B fusion protein construct. The fusion protein band is indicated in Figure 2 by the letter “A”. pET21 METTL7B was also identified by proteomics in both the purified protein fraction, and in the 55 kDa band in lane 1 of Figure 2 (Extended Data Tables 2 and 3). The lower bands around 30-35 kDa were determined by western blot to be co-purified affinity tag GST protein without METTL7B, as indicated by “B” and “C” in Figure 2. Tryptic digest showed that the remaining background bands are co-purified *E. coli* proteins (Extended Data Table 4).

**Figure 2.**
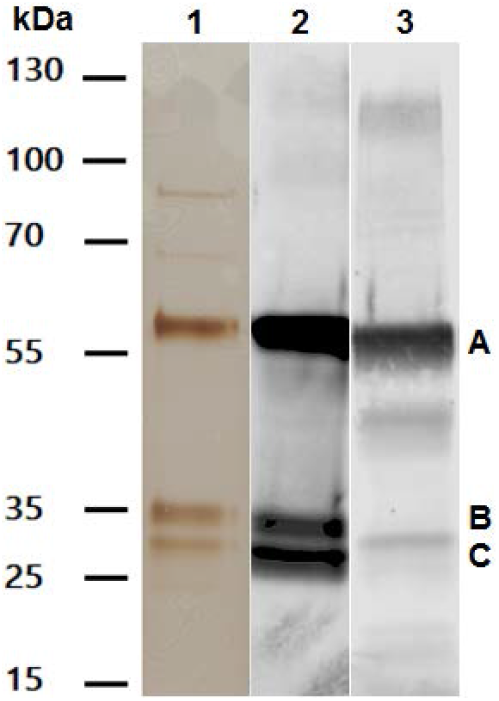
Analysis of purified pET21 METTL7B: **Lane 1)** SDS-PAGE silver stain of a representative gel showing purified pET21 METTL7B. The gel was loaded with a total of 1 μg total protein as determined by A_280_. **Lane 2)** anti-GST western blot of purified pET21 METTL7B. The gel was loaded with a total of 0.1 μg total protein. **Lane 3)** anti-METTL7B western blot of purified pET21 METTL7B. The gel was loaded with 0.1 μg total protein. Molecular weight markers from the PageRuler Plus Prestained Protein Ladder are shown on the left. The pET21 METTL7B band is marked by the letter A. The lower molecular bands, marked by letters B and C, are fusion protein fragments containing the dual His-GST affinity tag.

### Substrate Specificity Testing and Kinetic Analysis of pET21 METTL7B

A number of known methyltransferase substrates as well as endogenous thiol compounds were screened for methylation using recombinant pET21 METTL7B. Substrates were screened at concentrations at least three times higher than previously reported K_m_ values to ensure detection of methylation activity. We accounted for non-enzymatic methylation, which has been reported for some potential substrates, by including boiled enzyme and buffer-only controls and subtracting that turnover from the experimental samples. Qualitative screening results are presented below in Table 1. A subset of the semi-quantitative screening results is shown in Extended Data Figure 1. Only aliphatic thiol compounds show significant methylation signal above baseline.

**Table 1:**
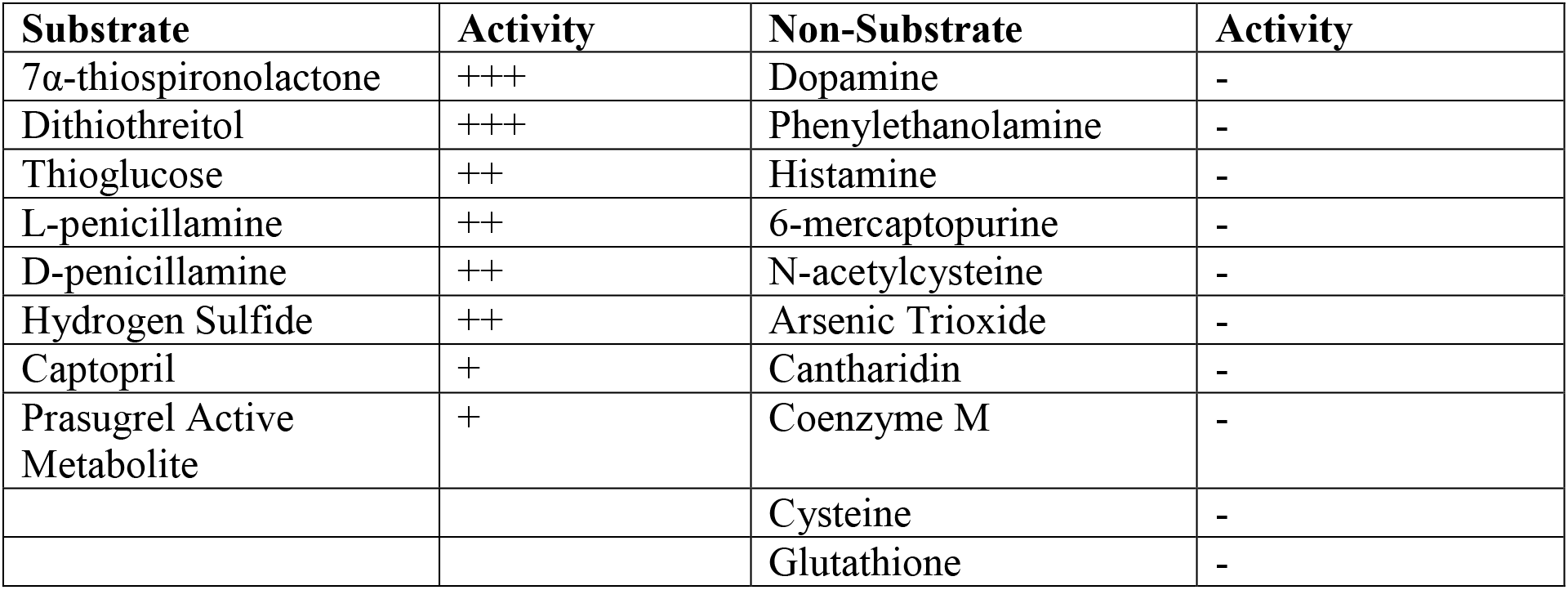
Relative Turnover of Probe Substrates with METTL7B.

We determined kinetic parameters for a subset of the identified substrates and they are presented below in Figure 3. All substrates were only methylated in the presence of SAM and catalytic activity was saturable and can be destroyed upon pre-boiling the enzyme. Additionally, kinetic experiments were conducted under conditions where methylation was linear with respect to incubation time and protein concentration (Extended Data Figures 2 and 3).

**Figure 3.**
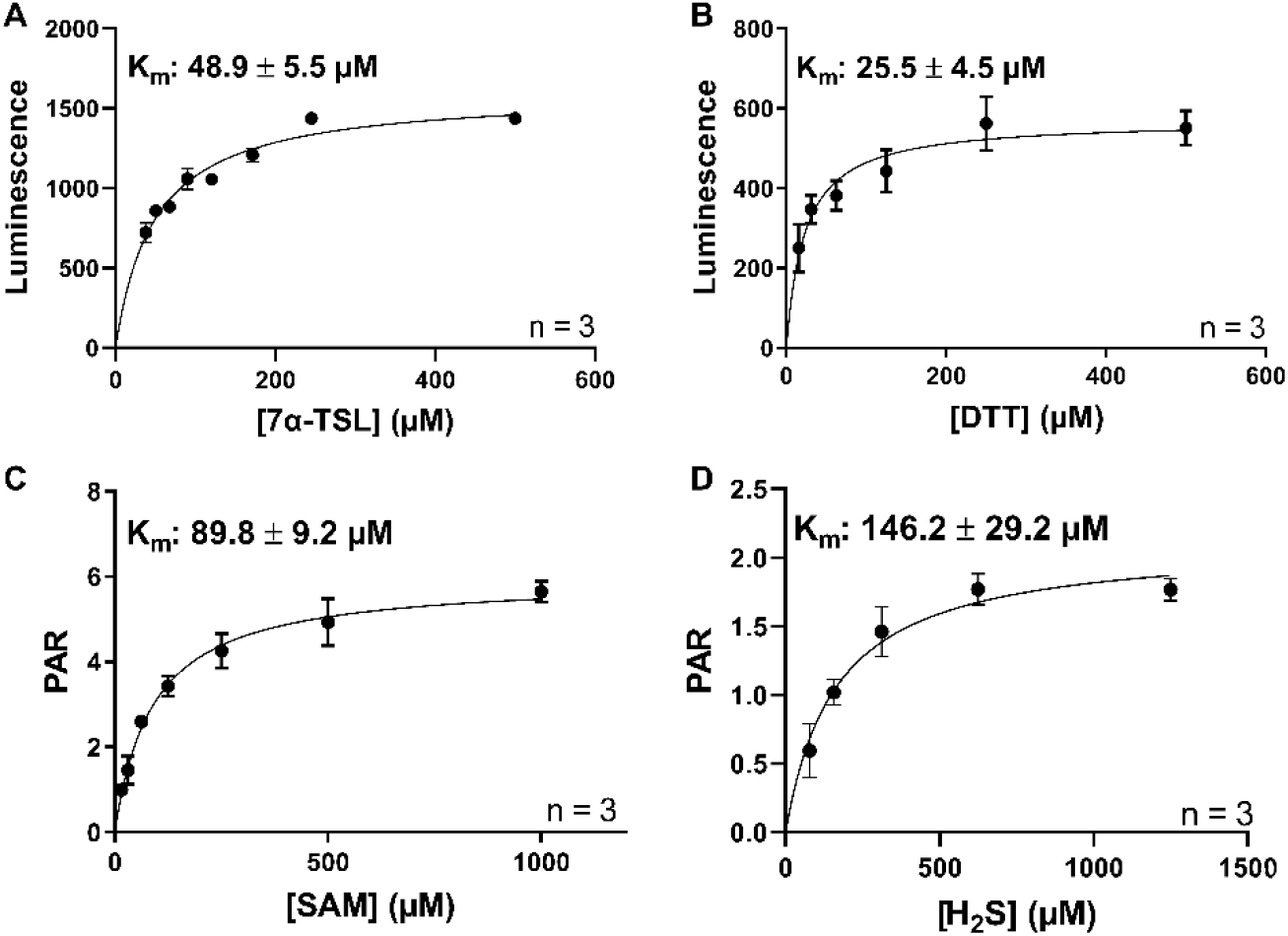
Rate of thiol methyl formation for pET21 METTL7B with multiple probe substrates: **A)** Luminescence vs concentration curves for 7α-thiospironolactone methylation as measured by SAH formation. **B)** Luminescence vs concentration curve for dithiothreitol methylation as measured by SAH formation. **C)** Peak area ratio (PAR) vs concentration curve for S-adenosyl-L-methionine use as measured by captopril methylation. **D)** PAR vs concentration curve for hydrogen sulfide methylation as measured by formation of methanethiol. All data is presented as the mean ± standard deviation of biological replicates.

Hydrogen sulfide and SAM kinetic curves were obtained using mass spectrometric methods measuring formation of methanethiol and *S*-methyl captopril respectively. 7α-thiospironolactone and dithiothreitol kinetic curves were obtained using the MTaseGlo kit (Promega) which measures the formation of S-adenosyl-L-homocysteine (SAH), the byproduct of all SAM-dependent methylation reactions.

Most substrates exhibit mid- to low-micromolar affinities to pET21 METTL7B. All substrates display classic Michaelis-Menten kinetics as evidenced by highly linear Eadie-Hofstee transformations of the data (Extended Data Figure 4).

## Discussion

The key finding in this paper is the METTL7B encodes for an alkyl thiol methyltransferase. We originally identified METTL7B as a candidate alkyl thiol methyltransferase by proteomic analysis of partially purified rat liver microsomes. Subsequent bioinformatics analysis determined that the human METTL7B had high sequence identity with the rat enzyme and has a putative SAM binding domain. We first manipulated *METTL7B* gene expression in human cell culture models to test that the gene product was associated with TMT activity using captopril as a probe substrate. We chose HepG2 and HeLa cells for gene knockdown and overexpression experiments because of their respectively high and low basal levels of *METTL7B* mRNA. Reduction of *METTL7B* gene, and protein, expression resulted in a decrease in captopril methylation. The opposite trend was observed upon gene overexpression, where increasing *METTL7B* gene and protein expression vastly increased captopril methylation.

We then designed a plasmid to express and purify recombinant full-length METTL7B to confirm that it was catalyzing the methylation event unequivocally. In our study, we discovered that glycerol greatly stabilized recombinant METTL7B in solution and that methylation activity was enhanced by adding dimyristoyl-*sn*-glycero-3-PG (DMPG) liposomes to reconstitute the protein. This was critical to maintain activity of an enzyme that is highly unstable which may have contributed to lack of characterization to date.

The METTL7B fusion protein catalyzes the *S*-methylation of multiple previously identified TMT-specific substrates in a SAM-dependent manner. We observed no methylation with a variety of probe substrates for other small molecule methyltransferases, as shown in Figure 3. The substrates that undergo methylation conform to the substrate specificity parameters previously determined using liver microsomes(1, 2, 24, 26). In general, METTL7B methylates compounds that contain an easily accessible aliphatic thiol functional group. It is important to note that METTL7B does not methylate 6-mercaptopurine, a classic thiopurine methyltransferase (TPMT) probe substrate(37). This further confirms that METTL7B catalyzes TMT-specific reactions rather than TPMT reactions. Additionally, consistent with prior reports, neither cysteine nor glutathione are substrates, but hydrogen sulfide is enzymatically methylated(23).

A potential key endogenous function of METTL7B is that it catalyzes the methylation of hydrogen sulfide to methanethiol which has been detected *in vivo* but the exact function and activity is still unknown. Maintenance of hydrogen sulfide homeostasis is crucial as it is known to play a large role in inflammatory processes, cell cycle, and cancer progression(38). In general, hydrogen sulfide exerts protective effects such as angiogenesis and cell growth at low concentrations. As hydrogen sulfide concentrations increase, its beneficial effects give way to toxicity, resulting in increased apoptosis(39, 40). Therefore, cancer cells with impaired hydrogen sulfide oxidation pathways, due to the hypoxic nature of tumors, likely rely heavily on methylation as a route of catabolism to prevent intracellular H_2_S levels from reaching toxic concentrations. Consequently, *METTL7B* is upregulated to potentially increase the rate of clearance of hydrogen sulfide or perhaps to increase formation of methanethiol. It is clear, however, that further research is required to fully characterize the role of METTL7B in the metabolism and homeostasis of hydrogen sulfide, especially in disease states that exhibit altered cellular redox states, such as the hypoxic interior of solid tumors. Additionally, it is important to investigate the role of methanethiol in cancer progression and its potential as a signaling molecule.

Overall, METTL7B possesses all of the known characteristics of the elusive human alkyl thiol methyltransferase (TMT) and should be renamed as alkyl thiol methyltransferase. Human METTL7B clearly catalyzes the SAM-dependent methyl transfer to exogenous and select endogenous thiol compounds, distinct from TPMT and other small molecule methyltransferases. METTL7B is involved in the metabolism of hydrogen sulfide, which may be important in cancer and inflammation where gene expression is highly upregulated and hydrogen sulfide levels are altered. Future work will focus on elucidating the *in vivo* role METTL7B plays in healthy and diseased tissue.

## Methods

### Materials

Mammalian overexpression plasmids and siRNA were purchased from Origene (Rockville, MD). HepG2 and HeLa cells were obtained from ATCC (Manassas, VA). Cell culture materials and lipofection reagents were purchased from ThermoFisher (Waltham, MA). Buffer salts were acquired from Sigma-Aldrich (St. Louis, MO) as well as methyltransferase probe substrates unless otherwise indicated. S-adenosyl-L-methionine and molecular biology kits were obtained from New England Biolabs (Ipswich, MA). Stellar Competent cells were purchased from Takara (Mountain View, CA). LOBSTR-BL21(DE3) competent cells were bought from Kerafast (Boston, MA). CHAPS detergent and UPLC-grade solvents were obtained from Fisher Scientific (Hampton, NH). Sequencing grade porcine trypsin and MTase-Glo Methyltransferase Assays were purchased from Promega (Madison, WI). 1,2-Dimyristoyl-*sn*-glycero-3-PG (DMPG) and mertansine were obtained from Cayman Chemical (Ann Arbor, MI). The active metabolite of prasugrel was a gift from Dr. Allan Rettie.

### HepG2 and HeLa Cell Culture

Cells were maintained and expanded using Dubelco’s Modified Eagle Medium supplemented with 10% fetal bovine serum and 0.1% penicillin/streptomycin. All cellular captopril methylation assays were conducted in serum-free media under optimized incubation conditions. Cells used for RNA isolation were washed with 1x phosphate buffered saline (PBS) prior to aspiration and storage at −80 °C until future use.

### Gene Expression Modulation

HepG2 cells were treated with Lipofectamine RNAiMax (Thermo Fisher Scientific, Waltham, MA) according to the manufacturer protocol, optimized for transfection duration. *GAPDH* gene knockdown using the Trilencer small interfering RNA (Origene, Rockville, MD) acted as the positive control for all gene expression knockdown experiments.

Cells were transfected in 12-well plates using a reverse transfection protocol. Briefly, *METTL7B* or scrambled siRNA was mixed with Lipofectamine RNAiMax in OptiMEM at room temperature for a final siRNA concentration of 50 nM. HepG2 cells were harvested using trypsin, pelleted, and resuspended to a final concentration of 200,000 cell/mL. Lipofectamine/siRNA stocks were added to culture plate wells, followed by 1 mL of cells, for a final concentration of 10 nM siRNA. Cells were allowed to incubate in the transfection media for 72 hours followed by RNA isolation or captopril methylation assays.

HeLa cells were treated with Lipofectamine 3000 (Thermo Fisher Scientific) according to the manufacturer protocol, optimized for transfection duration. Cells were transfected in 12-well plates via reverse transfection where purified empty or FLAG-tagged *METTL7B* overexpression plasmids (Origene) were mixed with P3000 reagent in OptiMEM at room temperature followed by Lipofectamine 3000. HeLa cells were harvested via trypsinization, pelleted, and resuspended to a final concentration of 200,000 cell/mL. Lipofectamine/plasmid stocks were added to culture plate wells, followed by 1 mL of cells, for a final plasmid concentration of 833 ng/mL. Cells were allowed to incubate in transfection media for 48 hours prior to RNA isolation or captopril methylation assays.

### Measurement of Gene Expression Changes

Cellular RNA was extracted using the MagMAX 96 Total RNA Isolation kit (Thermo Fisher Scientific) according to the manufacturer protocol. RNA quality (A_260_/A_280_) and concentration was assayed using a NanoDrop spectrophotometer. Isolated RNA was used to create cDNA using the High Capacity RNA-to-cDNA kit (Thermo Fisher Scientific) according to the manufacturer protocol. Subsequently, reverse-transcription polymerase chain reaction (RT-PCR) was conducted using an Applied Biosystems StepOnePlus Real-Time PCR System with TaqMan FAM reporter primers for *METTL7B, GAPDH,* and the housekeeping gene, *GusB*.

Expression level changes upon siRNA or plasmid treatment were determined using the ΔΔC_T_ method(41). In this method, *METTL7B* cycle threshold (C_t_) values are normalized to *GusB* C_t_ values in all samples, yielding a ΔC_T_ value. Relative gene expression changes are then calculated between treated and control cells using 2^−ΔΔCT^.

### Cellular Captopril Methylation Assay

Cells with altered *METTL7B* gene expression were created as described above. After the appropriate transfection period, cells were washed with 1x PBS and treated with serum-free media containing 500 μM captopril. Cell media aliquots were sampled after 24 hours and the S-methyl captopril metabolite was measured via liquid chromatography-tandem mass spectrometry (LC/MS-MS) and multiple reaction monitoring (MRM).

The LC-MS/MS system used for captopril methylation analysis was a Waters Xevo TQS mass spectrometer paired with a Waters Acquity LC. Compound separation was achieved using a 2.1×100 mm Ascentis Express RP Amide column and 0.1% formic acid in water and 0.1% formic acid in methanol as solvents A and B respectively. Column temperature was maintained at 50 °C at all times. Chromatographic separation was obtained using the following gradient: solvent B was held at 30% from 0 to 3 min, then held at 90% from 3 to 7 min, followed by re-equilibration to the starting conditions for another 3 min for a total run time of 10 min. Flow rate was held constant at 0.2 mL/min and flow was only diverted to the mass spectrometer between 2 to 7.5 min.

*S*-methyl captopril and the internal standard, d_3_-*S*-methyl captopril, were monitored in positive mode. The monitored mass transitions m/z+ were 232.1 > 89 and 232.1 > 116 (S-methyl captopril) as well as 235.1 > 91.9 and 235.1 > 115.9 (internal standard). The MS conditions were as follows: collision energy 15 V, cone voltage 30 V, capillary voltage 3.2 kV, desolvation temperature 450 °C, desolvation gas flow 1,000 L/hr and cone gas 150 L/hr.

### METTL7B Expression and Purification

Recombinant *METTL7B* was cloned in *E. coli* using a unique expression plasmid created in our lab. The expression plasmid backbone was obtained from pET21-10XHis-GST-HRV-dL5 which was a gift from Marcel Bruchez (Addgene plasmid # 73214; http://n2t.net/addgene:73214; RRID:Addgene_73214). The human *METTL7B* gene sequence was inserted into the plasmid using BamHI and EcoRI restriction sites and general molecular biology techniques. All plasmid inserts were sequenced by Eurofins Genomics and sequencing histograms were analyzed using FinchTV software.

Expression plasmids were propagated using heat-shocked Stellar cells. Individual colonies were used to create glycerol stocks and purify expression plasmids which were then sequenced for potential mutations. Validated plasmids were used to transform competent LOBSTR-BL21(DE3) *E. coli* via heat shock. Unless otherwise noted, all *E. coli* growth occurred on an orbital shaker at 250 rpm, 37 °C, and in the presence of 100 μg/mL ampicillin.

To express recombinant protein, LOBSTR-BL21(DE3) overnight cultures were added to ampicillin-containing TB expression media at a ratio of 1:100. Briefly, cells were grown for 3 hours under normal growth conditions. Then, METTL7B production was induced via addition of isopropyl ß-D-1-thiogalactopyranoside (IPTG) to a final concentration of 1 mM. The temperature was reduced to 15 °C and the cells were grown for an additional 24 hours. Cells were harvested via gentle centrifugation and the collected pellets were stored at −80 °C until future processing.

Frozen cell pellets were thawed on ice in a 4 °C cold cabinet overnight prior to lysis. Lysis was conducted by resuspending the cell pellet in cell lysis buffer (50 mM KPi pH 7.0, 20% glycerol, 150 mM NaCl, 10 mM CHAPS, EDTA-free Halt Protease Inhibitor Cocktail) supplemented with 100 μg/mL lysozyme (Sigma Aldrich). The cell solution was rotated end-over-end at 4 °C until the solution had become extremely viscous. Then, the cell lysate was treated with 100 μg/mL DNA Nuclease I (Sigma Aldrich) and rotated at 4 °C or until no longer viscous. The lysate was then centrifuged at 48,000 g for 30 minutes at 4 °C and the resulting supernatant was retained for subsequent purification steps.

Purification was conducted using the ÄKTA start chromatography system (GE Healthcare). Cell lysate supernatant was applied to a pre-packed and conditioned HisPur Ni-NTA column (ThermoFisher) overnight at a low flow rate (0.5 mL/min). The column was subsequently washed with Ni-NTA purification buffer (50 mM KPi pH 7.0, 20% glycerol, 10 mM CHAPS, 300 mM NaCl) containing 50 mM imidazole until A_280_ readings stabilize. Protein was eluted from the column with purification buffer containing 300 mM imidazole until A_280_ readings stabilized.

The HisPur Ni-NTA column eluent was directly applied to a pre-conditioned GSTrapFF column at a flow rate of 1 mL/min for 4 hours. The column was then washed with GSTrapFF purification buffer (50 mM KPi pH 7.0, 20% glycerol, 10 mM CHAPS, 150 mM NaCl) until A_280_ had decreased to baseline. Recombinant protein was eluted from the column using purification buffer containing 10 mM reduced glutathione and adjusted to pH 8.0. Pooled eluent was concentrated to appropriate working concentrations using Amicon Centriprep 10K molecular weight cutoff centrifugal filter units. Final protein concentration was determined by A280 measurement and stocks were aliquoted and stored at −80 °C until future use.

### In vitro Captopril Methylation Using Recombinant METTL7B

*In vitro* captopril methylation was conducted using purified METTL7B fusion protein. The reaction buffer (50 mM KPi pH 7.0, 10 mM CHAPS, 20% glycerol, 150 mM NaCl, and 9 mg/mL DMPG) was placed in a sonication water bath until the solution was clear to help form DMPG liposomes. Recombinant enzyme was added at a ratio of 85:1 DMPG:METTL7B and allowed to incubate on ice for 30 minutes. Following the addition of captopril, the enzyme was pre-equilibrated at 37 °C for 2 minutes before initiation by addition of SAM to a reaction volume of 150 μL. The final concentration of captopril was varied to collect the kinetic information and the final concentration of SAM was held at 750 μM. The reaction was incubated for 25 minutes and then quenched via addition of 15% (w/v) zinc sulfate in a 1:5 dilution to total reaction volume. The quenched solution was incubated on ice for 10 minutes followed by a 1:6 addition of a saturated barium hydroxide solution containing the d_3_-S-methyl captopril internal standard. Following a second 10-minute incubation on ice, the solution was centrifuged at 5,000 xg for 15 minutes to pellet all precipitated proteins and salts.

Following centrifugation, 75 μL of supernatant was transferred to an opaque polypropylene strip-well tube containing 5 μL of 2 M sodium hydroxide. Unreacted captopril was derivatized at room temperature for 1 hour in the dark via addition of 20 μL of 2.5 M maleimide to reduce ion suppression from non-methylated captopril. Derivatized samples were centrifuged and the supernatant was analyzed by LC-MS/MS as previously described.

### In vitro Hydrogen Sulfide Methylation Using Recombinant METTL7B

Protein concentration and incubation with DMPG liposomes was conducted the same as described above. All steps were conducted in a glove box under nitrogen unless otherwise indicated. Recombinant enzyme was aliquoted into a polypropylene deep-well plate on ice along with SAM and NaSH for a final volume of 150 μL and 0.09 mg/mL and 83.3 μM for protein and SAM concentrations respectively. The plate was capped with a silicon mat and placed in a 37 °C water bath for 45 min under normal atmosphere. After incubation, the plate was placed back on ice under nitrogen and quenched via a 1:15 addition of 0.3 M sodium hydroxide. 110 μL of the quenched reaction solution was added to 50 μL of 20 mM monobromobimane (MBB), based off of published H_2_S derivatization method(42–44). Once capped under nitrogen, the reaction plate was incubated at room temperature on an orbital shaker at 450 rpm for 30 min.

The MBB derivatization was quenched by addition of 50 μL of 200 mM 5-sulfosalicylic acid and 10 μL of the ethyl 2-aminothiazole carboxylate (EATC) internal standard. Protein was precipitated by addition of 15% (w/v) zinc sulfate and barium hydroxide as previously detailed. Samples were centrifuged at 4,000 xg for 15 min and the supernatant was analyzed by LC-MS/MS.

The LC-MS/MS system used for hydrogen sulfide methylation analysis was a Waters Xevo TQS mass spectrometer paired with a Waters Acquity LC. Compound separation was achieved using a 2.1×150 mm Acquity UPLC BEH Shield RP column and 0.2% acetic acid in water and 0.2% acetic acid in acetonitrile as solvents A and B respectively. Column temperature was maintained at 25 °C at all times. Chromatographic separation was obtained using the following gradient: solvent B was held at 40% from 0 to 1 minutes, ramped to 90% from 1 to 3.5 minutes, held at 90% from 4.5 to 5 minutes followed by re-equilibration to the starting conditions for another minute. Flow rate was held constant at 0.3 mL/min and flow was only diverted to the mass spectrometer from 1 to 4.5 minutes.

Derivatized methanethiol and the internal standard, EATC, were monitored in positive mode. The monitored mass transitions m/z+ were 239.22 > 175.24 and 239.22 > 192.2 (derivatized methanethiol) as well as 173.17 > 72.11 and 173.17 > 127.06 (internal standard). The MS conditions were as follows: collision energy 24, 10, 24, 16 V for each transition respectively, cone voltage 56 V, capillary voltage 2.9 kV, desolvation temperature 450 °C, desolvation gas flow 1,000 L/hr and cone gas 150 L/hr.

### Protein Purity Analysis

All SDS-PAGE silver stain analysis was conducted using NuPAGE 4-12% Bis-Tris gels in the XCell SureLock Mini-Cell Electrophoresis system using PageRuler Plus Prestained Protein Ladder as a molecular weight marker. Samples were prepared using NuPAGE LDS Sample Buffer sample buffer and 1.4 M ß-mercaptoethanol before boiling for 5 minutes. Gels were run at room temperature, at a constant 200 volts, and developed using previously published silver staining protocols(45).

All western blot analyses were conducted using the XCell SureLock Mini-Cell Electrophoresis system, PageRuler Plus Prestained Protein Ladder, and NuPAGE 10-20% Tricine gels. Samples were prepared using 4X Protein Loading Buffer (LiCor) and 0.7 M ß-mercaptoethanol before boiling for 5 minutes. After initial SDS-PAGE separation, the gel was removed from the cassette and placed with PVDF blotting membrane into the XCell II Blot Module according to the manufacturer protocol. Blot transfer was conducted over 1 hour at a constant 30 volts on ice. The blot was blocked using Odyssey Blocking Buffer (LiCor) for 1 hour at room temperature. A primary antibody incubation was conducted overnight using the suggested dilution factor for the rabbit anti-METTL7B (Sigma Aldrich), anti-FLAG (Cell Signaling), anti-GST (Cell Signaling), or anti-ß actin (Cell Signaling) antibodies. The secondary antibody incubation was conducted for 1 hour at room temperature using IRDye 680RD goat anti-rabbit antibody (LiCor). Western blots were scanned using an Odyssey gel scanner. Blot images were visualized using Image Studio Version 4.0 software.

### Tryptic Digest

In-gel tryptic digests of silver stained SDS-PAGE gels were conducted following the method published by Shevchenko(45). Briefly, the protein band was excised from the gel and dehydrated with neat acetonitrile. Protein bands were then treated with 10 mM dithiothreitol (DTT) solution and incubated at 56 °C to reduce all proteins. The reduced bands were treated with 55 mM iodoacetamide at room temperature in the dark to alkylate all exposed cysteine side chains. Finally, the bands were incubated overnight at 37 °C with 13 ng/μL trypsin-containing solution. Tryptic digestion peptides were extracted from the gel bands the following day and concentrated in a centrifugal evaporator. Concentrated peptides were analyzed using a high-resolution mass spectrometer, a Finnigan LTQ Orbitrap in our case, and then used to identify the protein of interest via ProteinProspector.

The LC-MS system used for proteomic analysis was a Finnigan LTQ Orbitrap coupled to a Waters Acquity LC. Peptides were separated using a 1×150 mm Acquity UPLC CSH C18 column and 0.1% formic acid in water and 0.1% formic acid in acetonitrile as solvents A and B. Separation was achieved using the following gradient: solvent B was held at 5% for the first two minutes, increased to 40% over the next 90 min, increased to 90% over the next five minutes and held for an additional 8 minutes, then re-equilibrated over five minutes. The flow rate was held at 0.06 mL/min and flow as diverted to the mass spectrometer from 2 to 95 minutes.

Peptides were analyzed using a data dependent scan method in positive mode. The initial high resolution scan from 300-2,000 m/z was conducted in the FTMS with 60,000 resolution. Four dependent scans were completed in the ion trap to obtain fragmentation. Dynamic exclusion was enabled which excluded the top 25 most intense ions after they had been selected twice over a four second window. The following mass spectrometer settings were used: sheath gas flow rate was 12 arb, spray voltage was 3.5 kV, capillary temperature was 350 °C, capillary voltage was 22 V, and tube lens voltage was 100 V.

### Substrate Screening

Substrate screening was primarily conducted using the MTase-Glo Assay (Promega). Briefly, recombinant METTL7B was prepared the same way as for use in the captopril methylation assay. SAM was added to the METTL7B protein stock for a final concentration of 50 μM and was aliquoted into a 384-well plate. Substrate was added to each well and the plate was covered using Parafilm before incubating at 37 °C for 1 hour. The incubation was quenched by a 1:5 addition of 0.5% trifluoro acetic acid. The samples were processed according to the manufacturer protocol and luminescence was recorded for each well using a Synergy HTX Multi-Mode Reader (BioTek).

### Data Analysis

All experiments were conducted with biological triplicates, and repeated at least two times on two different days. All data is reported as the mean ± standard deviation, however individual data points from multiple experiments are presented when possible. Statistical significance was determined by a two-tailed unpaired *t* test with a threshold *P* value of 0.05. Kinetic parameter Km values were obtained through non-linear regression analysis using GraphPad Prism, version 8.3.1 for Windows (GraphPad Software, La Jolla, CA).

## Data Availability Statement

The proteomic data that support the findings of this study are available from PeptideAtlas, tagged as “pET21METTL7B”. All other data are available from the corresponding author upon reasonable request.

## Materials Availability Statement

Unique materials used when conducting the experiments detailed in this study are available from the corresponding author upon reasonable request.

