## Supplemental Tables 1-4 and Supplemental Figures 1-4 for "Human *METTL7B* Encodes an Alkyl Thiol Methyltransferase that Methylates Hydrogen Sulfide"

**Supplemental Data:**

**Supplemental Data Table 1: Proteins Identified in TMT-active RLM Purification Fraction**

| Rank | Acc # | 11_12_2015_005/DEAE Post-Void FT | | | Protein MW | Species | Protein Name |
|---|---|---|---|---|---|---|---|
| | | Num Unique | % Cov | Best Expect Val | | | |
| 1 | P10634 | 20 | 46.2 | 2.0e-10 | 56684.4 | RAT | Cytochrome P450 2D26 |
| 2 | P06761 | 19 | 34.1 | 2.4e-10 | 72347.6 | RAT | 78 kDa glucose-regulated protein |
| 3 | P18163 | 18 | 37.5 | 2.7e-8 | 78179.4 | RAT | Long-chain-fatty-acid--CoA ligase 1 |
| 4 | P14141 | 14 | 85.8 | 1.8e-8 | 29431.6 | RAT | Carbonic anhydrase 3 |
| 5 | P07687 | 18 | 49.0 | 2.4e-7 | 52582.0 | RAT | Epoxide hydrolase 1 |
| 6 | P08683 | 14 | 39.0 | 3.9e-9 | 57181.6 | RAT | Cytochrome P450 2C11 |
| 7 | Q64573 | 11 | 31.6 | 1.5e-10 | 62309.0 | RAT | Liver carboxylesterase 4 |
| 8 | P46462 | 14 | 30.5 | 1.0e-7 | 89349.6 | RAT | Transitional endoplasmic reticulum ATPase |
| 9 | P11442 | 10 | 9.4 | 1.7e-8 | 191600.4 | RAT | Clathrin heavy chain 1 |
| 10 | Q63081 | 9 | 34.3 | 8.4e-9 | 48173.8 | RAT | Protein disulfide-isomerase A6 |
| 11 | P02692 | 8 | 74.0 | 2.7e-9 | 14272.7 | RAT | Fatty acid-binding protein, liver |
| 12 | P08541 | 10 | 24.5 | 3.6e-7 | 60986.1 | RAT | UDP-glucuronosyltransferase 2B2 |
| 13 | P08010 | 10 | 54.1 | 3.2e-7 | 25702.9 | RAT | Glutathione S-transferase Mu 2 |
| 14 | P04905 | 9 | 51.8 | 2.1e-6 | 25914.2 | RAT | Glutathione S-transferase Mu 1 |
| 15 | P30839 | 12 | 30.0 | 2.9e-7 | 54082.1 | RAT | Fatty aldehyde dehydrogenase |
| 16 | P00173 | 8 | 56.7 | 2.8e-8 | 15355.3 | RAT | Cytochrome b5 |
| 17 | P10719 | 6 | 19.8 | 3.4e-8 | 56354.0 | RAT | ATP synthase subunit beta, mitochondrial |
| 18 | Q9EQ76 | 6 | 14.3 | 1.1e-8 | 59960.9 | RAT | Dimethylaniline monooxygenase [N-oxide-forming] 3 |
| 19 | Q66HD0 | 7 | 13.4 | 4.7e-8 | 92771.7 | RAT | Endoplasmin |
| 20 | P52759 | 5 | 60.6 | 7.0e-10 | 14303.6 | RAT | 2-iminobutanoate/2-iminopropanoate deaminase |
| 21 | P29147 | 6 | 23.3 | 4.2e-8 | 38202.2 | RAT | D-beta-hydroxybutyrate dehydrogenase, mitochondrial |
| 22 | P55006 | 6 | 23.7 | 6.1e-11 | 35737.0 | RAT | Retinol dehydrogenase 7 |
| 23 | Q562C4 | 6 | 36.5 | 2.1e-6 | 27904.1 | RAT | Methyltransferase-like protein 7B |
| 24 | P04797 | 6 | 27.9 | 2.3e-10 | 35828.3 | RAT | Glyceraldehyde-3-phosphate dehydrogenase |
| 25 | P15999 | 7 | 20.1 | 2.5e-8 | 59754.1 | RAT | ATP synthase subunit alpha, mitochondrial |
| 26 | P08011 | 4 | 23.9 | 6.6e-9 | 17471.7 | RAT | Microsomal glutathione S-transferase 1 |
| 27 | P60711 | 4 | 17.1 | 8.9e-8 | 41737.1 | RAT | Actin, cytoplasmic 1 |
| 28 | Q9Z0V5 | 6 | 34.8 | 3.7e-6 | 31007.7 | RAT | Peroxiredoxin-4 |
| 29 | P70580 | 3 | 19.5 | 7.0e-9 | 21598.2 | RAT | Membrane-associated progesterone receptor component 1 |
| 30 | P27364 | 5 | 16.6 | 4.3e-6 | 42206.6 | RAT | 3 beta-hydroxysteroid dehydrogenase type 5 |
| 31 | P00502 | 4 | 22.5 | 2.0e-8 | 25607.3 | RAT | Glutathione S-transferase alpha-1 |
| 32 | Q6UPE0 | 4 | 12.5 | 3.6e-7 | 66389.1 | RAT | Choline dehydrogenase, mitochondrial |
| 33 | P00388 | 4 | 10.0 | 5.3e-7 | 76963.5 | RAT | NADPH--cytochrome P450 reductase |
| 34 | P02091 | 4 | 25.2 | 9.1e-7 | 15979.5 | RAT | Hemoglobin subunit beta-1 |
| 35 | P16232 | 5 | 15.6 | 7.8e-7 | 31883.6 | RAT | Corticosteroid 11-beta-dehydrogenase isozyme 1 |
| 36 | P05182 | 4 | 11.0 | 3.9e-6 | 56627.4 | RAT | Cytochrome P450 2E1 |
| 37 | P10867 | 3 | 9.3 | 2.8e-6 | 50615.8 | RAT | L-gulonolactone oxidase |

| | | | | | | | |
|---|---|---|---|---|---|---|---|
| 38 | Q6UPE1 | 3 | 7.0 | 5.6e-8 | 68198.7 | RAT | Electron transfer flavoprotein-ubiquinone oxidoreductase, mitochondrial |
| 39 | Q09073 | 3 | 16.8 | 4.5e-7 | 32901.6 | RAT | ADP/ATP translocase 2 |
| 40 | P27867 | 2 | 10.1 | 1.4e-7 | 38234.9 | RAT | Sorbitol dehydrogenase |
| 41 | P11030 | 3 | 41.4 | 4.2e-7 | 10027.5 | RAT | Acyl-CoA-binding protein |
| 42 | P55159 | 3 | 13.8 | 5.7e-6 | 39358.3 | RAT | Serum paraoxonase/arylesterase 1 |
| 43 | Q9ES38 | 4 | 9.3 | 1.6e-6 | 76266.3 | RAT | Bile acyl-CoA synthetase |
| 44 | Q68FP2 | 2 | 7.9 | 2.1e-6 | 39458.5 | RAT | Serum paraoxonase/lactonase 3 |
| 45 | P01946 | 3 | 37.3 | 1.0e-6 | 15328.7 | RAT | Hemoglobin subunit alpha-1/2 |
| 46 | Q5PPL3 | 1 | 5.2 | 3.5e-8 | 40412.0 | RAT | Sterol-4-alpha-carboxylate 3-dehydrogenase, decarboxylating |
| 47 | P20070 | 2 | 12.0 | 4.9e-7 | 34174.9 | RAT | NADH-cytochrome b5 reductase 3 |
| 48 | Q62730 | 1 | 4.5 | 5.5e-8 | 41967.5 | RAT | Estradiol 17-beta-dehydrogenase 2 |
| 49 | P13107 | 2 | 5.1 | 7.4e-7 | 56384.9 | RAT | Cytochrome P450 2B3 |
| 50 | P04642 | 2 | 8.7 | 3.8e-6 | 36450.8 | RAT | L-lactate dehydrogenase A chain |
| 51 | P20816 | 2 | 5.0 | 4.6e-5 | 57969.5 | RAT | Cytochrome P450 4A2 |
| 52 | Q6AYS8 | 1 | 5.4 | 7.7e-8 | 32938.0 | RAT | Estradiol 17-beta-dehydrogenase 11 |
| 53 | P36365 | 3 | 8.1 | 3.8e-5 | 59825.9 | RAT | Dimethylaniline monooxygenase [N-oxide-forming] 1 |
| 54 | P24470 | 1 | 3.6 | 9.9e-7 | 56433.5 | RAT | Cytochrome P450 2C23 |
| 55 | P11240 | 2 | 21.9 | 6.2e-6 | 16129.7 | RAT | Cytochrome c oxidase subunit 5A, mitochondrial |
| 56 | Q63276 | 1 | 4.0 | 1.9e-7 | 46465.1 | RAT | Bile acid-CoA:amino acid N-acyltransferase |
| 57 | Q5XI60 | 1 | 6.2 | 1.5e-5 | 23313.3 | RAT | Receptor expression-enhancing protein 6 |
| 58 | P07632 | 1 | 8.4 | 4.1e-6 | 15911.8 | RAT | Superoxide dismutase [Cu-Zn] |
| 59 | Q0ZHH6 | 1 | 3.3 | 1.9e-6 | 60586.5 | RAT | Atlastin-3 |
| 60 | Q02253 | 1 | 3.2 | 5.9e-6 | 57808.1 | RAT | Methylmalonate-semialdehyde dehydrogenase [acylating], mitochondrial |
| 61 | P81828 | 1 | 13.9 | 3.2e-6 | 11068.0 | RAT | Urinary protein 2 |
| 62 | P05179 | 2 | 6.7 | 6.6e-5 | 56187.6 | RAT | Cytochrome P450 2C7 |
| 63 | P05183 | 1 | 2.6 | 8.5e-6 | 57732.2 | RAT | Cytochrome P450 3A2 |
| 64 | P51635 | 1 | 4.0 | 2.4e-6 | 36506.2 | RAT | Alcohol dehydrogenase [NADP(+)] |
| 65 | P00884 | 1 | 4.1 | 2.3e-6 | 39618.5 | RAT | Fructose-bisphosphate aldolase B |
| 66 | P02706 | 2 | 7.4 | 7.3e-5 | 32849.2 | RAT | Asialoglycoprotein receptor 1 |
| 67 | Q8K4C0 | 2 | 5.3 | 2.8e-5 | 60056.1 | RAT | Dimethylaniline monooxygenase [N-oxide-forming] 5 |
| 68 | Q5M9I5 | 1 | 20.2 | 7.5e-6 | 10423.7 | RAT | Cytochrome b-c1 complex subunit 6, mitochondrial |
| 69 | P19225 | 2 | 4.1 | 2.6e-5 | 56157.2 | RAT | Cytochrome P450 2C70 |
| 70 | P97524 | 1 | 1.8 | 4.5e-5 | 70694.5 | RAT | Very long-chain acyl-CoA synthetase |
| 71 | P50137 | 1 | 2.7 | 2.8e-5 | 67644.2 | RAT | Transketolase |
| 72 | Q5I0E7 | 1 | 7.2 | 7.5e-5 | 27028.3 | RAT | Transmembrane emp24 domain-containing protein 9 |
| 73 | P81827 | 1 | 10.9 | 8.2e-5 | 10960.0 | RAT | Urinary protein 1 |
| 74 | Q9JJ46 | 1 | 6.1 | 1.7e-4 | 26737.8 | RAT | 3-beta-hydroxysteroid-Delta(8),Delta(7)-isomerase |

### Supplemental Data Table 2: Human Peptides Identified from Purified pET21 METTL7B

1 **Acc. #:** Q6UX53 **Uniprot ID:** MET7B_HUMAN **Species:** HUMAN **Name:** Methyltransferase-like protein 7B
**Organism:** Homo sapiens **Gene:** METTL7B **Existence:** Evidence at protein level **Version:** 2
**Protein MW:** 27775.1 **Protein pI:** 8.7 **Protein Length:** 244 **Index:** 234488

| Num Unique | % Cov | Best Disc Score | Best Expect Val |
|---|---|---|---|
| 10 | 63.1 | 5.16 | 5.9e-9 |

| m/z | z | ppm | DB Peptide | Score | Expect | # in DB |
|---|---|---|---|---|---|---|
| 1214.0861 | 2 | 2.7 | VALLELGCGTGANFQFYPPGCR | 49.8 | 5.9e-9 | 1 |
| 855.3956 | 2 | 2.3 | DLENAQFSEIQMER | 45.0 | 1.3e-8 | 1 |
| 1127.5265 | 2 | 0.55 | ETWKDLENAQFSEIQMER | 52.1 | 1.1e-7 | 1 |
| 728.8484 | 2 | 0.41 | VTCLDPNPHFEK | 47.6 | 1.4e-7 | 1 |
| 765.4084 | 2 | 1.5 | SYFPYLMAVLTPK | 39.8 | 5.6e-7 | 1 |
| 594.7665 | 2 | 0.75 | HIGDGCCLTR | 31.6 | 2.3e-6 | 1 |
| 1071.0376 | 4 | -0.80 | RVLRPGGVLFFWEHVAEPYGSWAFMWQQVFEPTWK | 30.3 | 5.5e-6 | 1 |
| 845.4191 | 3 | 0.63 | QLADGSMDVVVCTLVLCSVQSPR | 31.7 | 5.8e-6 | 1 |
| 560.7768 | 2 | 0.68 | FVVAPGEDMR | 31.6 | 1.7e-5 | 1 |
| 617.8421 | 2 | 0.33 | WLPVGPHIMGK | 26.5 | 2.0e-5 | 1 |

### Supplemental Data Table 3: METTL7B Peptides Identified from SDS-PAGE In-Gel Digest

1 **Acc. #:** Q6UX53 **Uniprot ID:** MET7B_HUMAN **Species:** HUMAN **Name:** Methyltransferase-like protein 7B
**Organism:** Homo sapiens **Gene:** METTL7B **Existence:** Evidence at protein level **Version:** 2
**Protein MW:** 27775.1 **Protein pI:** 8.7 **Protein Length:** 244 **Index:** 234488

```
1   MDILVPLLQL LVLLLTLPLH LMALLGCWQP LCKSYFPYLM AVLTPKSNRK MESKKRELFS QIKGLTGASG KVALLELGCG
81  TGANFQFYPP GCRVTCLDPN PHFEKFLTKS MAENRHLQYE RFVVAPGEDM RQLADGSMDV VVCTLVLCSV QSPRKVLQEV
161 RRVLRPGGVL FFWEHVAEPY GSWAFMWQQV FEPTWKHIGD GCCLTRETWK DLENAQFSEI QMERQPPPLK WLPVGPHIMG
241 KAVK
```

| Num Unique | % Cov | Best Disc Score | Best Expect Val |
|---|---|---|---|
| 6 | 27.9 | 3.47 | 3.4e-6 |

| m/z | z | ppm | Score | Expect | # in DB |
|---|---|---|---|---|---|
| 828.7350 | 3 | 4.3 | 30.9 | 3.4e-6 | 1 |
| 891.9000 | 2 | -2.0 | 28.9 | 7.7e-6 | 1 |
| 757.3594 | 2 | 0.75 | 38.0 | 1.9e-5 | 1 |
| 883.9045 | 2 | 0.19 | 34.8 | 3.1e-5 | 1 |
| 589.2875 | 2 | 0.60 | 28.9 | 1.4e-4 | 1 |
| 623.2773 | 2 | 0.82 | 16.4 | 1.9e-4 | 1 |

**Supplemental Data Table 4: *E. coli* Proteins Identified Alongside Purified pET21 METTL7B**

| Rank | Acc # | 1_11_2019_001a/1_11_19 pET21 METTL7B GSTrapFF Eluent E coli 1 | | | | Protein MW | Species | Protein Name |
|---|---|---|---|---|---|---|---|---|
| | | Num Unique | % Cov | Best Disc Score | Best Expect Val | | | |
| 1 | P0A6Y8 | 5 | 11.6 | 4.67 | 4.8e-8 | 69115.5 | ECOLI | Chaperone protein DnaK |
| 2 | P0A6N2 | 4 | 12.7 | 2.78 | 1.6e-5 | 43314.0 | ECOL6 | Elongation factor Tu |
| 3 | P0A7L0 | 3 | 17.9 | 3.61 | 2.7e-6 | 24729.8 | ECOLI | 50S ribosomal protein L1 |
| 4 | P02413 | 2 | 18.1 | 4.82 | 2.5e-8 | 14980.5 | ECOLI | 50S ribosomal protein L15 |
| 5 | P0A7J3 | 3 | 23.6 | 2.76 | 1.2e-5 | 17711.7 | ECOLI | 50S ribosomal protein L10 |
| 6 | P0A7J7 | 2 | 16.2 | 4.32 | 2.1e-7 | 14875.5 | ECOLI | 50S ribosomal protein L11 |
| 7 | P0A7K2 | 2 | 19.0 | 3.83 | 1.8e-6 | 12295.3 | ECOLI | 50S ribosomal protein L7/L12 |
| 8 | P60723 | 3 | 15.9 | 3.76 | 6.3e-8 | 22086.7 | ECOLI | 50S ribosomal protein L4 |
| 9 | Q1RGI7 | 2 | 8.8 | 3.58 | 3.6e-7 | 41044.7 | ECOUT | Chaperone protein DnaJ |
| 10 | P0A7R5 | 2 | 24.3 | 3.93 | 5.9e-7 | 11735.7 | ECOLI | 30S ribosomal protein S10 |
| 11 | P0C054 | 2 | 15.3 | 2.85 | 1.7e-5 | 15773.8 | ECOLI | Small heat shock protein IbpA |
| 12 | P60422 | 3 | 16.8 | 3.13 | 9.5e-7 | 29860.7 | ECOLI | 50S ribosomal protein L2 |
| 13 | P0A7W1 | 1 | 11.4 | 5.30 | 3.3e-9 | 17603.5 | ECOLI | 30S ribosomal protein S5 |
| 14 | P62399 | 3 | 22.3 | 2.56 | 1.7e-6 | 20301.8 | ECOLI | 50S ribosomal protein L5 |
| 15 | P0ADB7 | 1 | 39.6 | 4.80 | 2.8e-8 | 4809.6 | ECOLI | Entericidin B |
| 16 | P0A7S9 | 2 | 26.3 | 2.74 | 8.6e-6 | 13099.5 | ECOLI | 30S ribosomal protein S13 |
| 17 | P0A7L3 | 2 | 16.1 | 2.62 | 4.0e-6 | 13497.1 | ECOLI | 50S ribosomal protein L20 |
| 18 | P0A7W7 | 1 | 10.8 | 4.05 | 8.0e-8 | 14126.7 | ECOLI | 30S ribosomal protein S8 |
| 19 | P0A7L8 | 1 | 16.5 | 3.41 | 3.4e-6 | 9124.5 | ECOLI | 50S ribosomal protein L27 |
| 20 | P0A7V3 | 1 | 5.2 | 3.35 | 7.5e-6 | 25983.4 | ECOLI | 30S ribosomal protein S3 |
| 21 | P0C058 | 2 | 12.7 | 2.04 | 3.5e-5 | 16093.3 | ECOLI | Small heat shock protein IbpB |
| 22 | P62593 | 2 | 8.0 | 1.43 | 2.0e-4 | 31515.5 | ECOLX | Beta-lactamase TEM |
| 23 | P02359 | 3 | 14.0 | 1.69 | 4.0e-5 | 20019.3 | ECOLI | 30S ribosomal protein S7 |
| 24 | P0AA10 | 1 | 7.0 | 2.18 | 1.8e-6 | 16018.7 | ECOLI | 50S ribosomal protein L13 |
| 25 | P60438 | 1 | 4.8 | 2.04 | 6.6e-5 | 22243.7 | ECOLI | 50S ribosomal protein L3 |
| 26 | P0A7V8 | 1 | 4.4 | 1.90 | 1.2e-4 | 23469.3 | ECOLI | 30S ribosomal protein S4 |
| 27 | P60624 | 1 | 9.6 | 1.43 | 4.4e-5 | 11316.3 | ECOLI | 50S ribosomal protein L24 |
| 28 | P0A9X9 | 1 | 21.4 | 1.01 | 2.0e-4 | 7403.3 | ECOLI | Cold shock protein CspA |
| 29 | P0A6A8 | 1 | 11.5 | 0.30 | 0.0038 | 8639.6 | ECOLI | Acyl carrier protein |

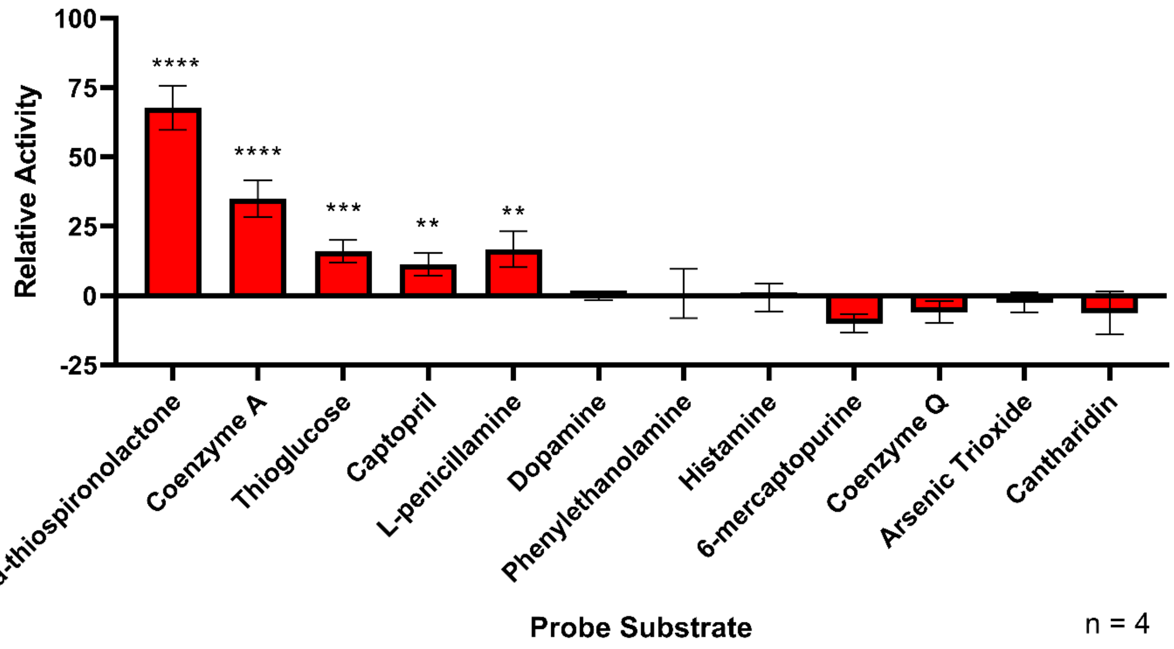


**Supplemental Data Figure 1: Semi-quantitative Screening of Select Methyltransferase Probe Substrates.** Multiple methyltransferase probe substrates were incubated for 1 hour with pET21 METTL7B at concentrations predicted to promote turnover. Formation of S-adenosyl-L-homocysteine was measured using the Promega MTaseGlo kit. Activity was normalized to dopamine, a catechol O-methyltransferase probe. All data is presented as the mean ± standard deviation. Significance was determined using unpaired two-tailed *t* test. ****$P<0.0001$. ***$P<0.001$. **$P<0.01$

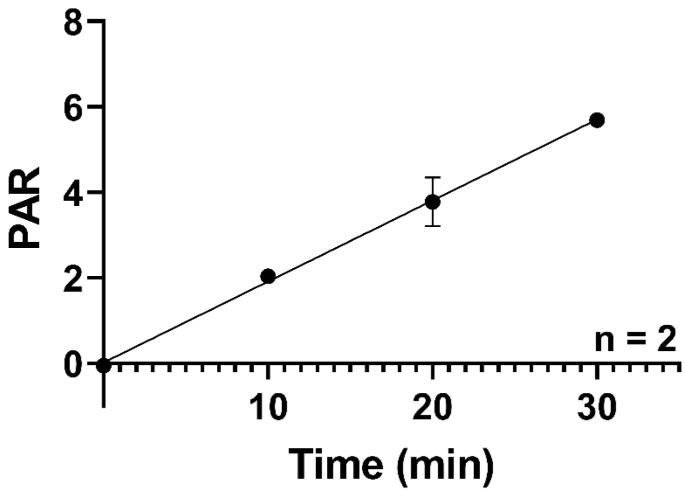


**Supplemental Data Figure 2: Time Linearity of pET21 METTL7B Captopril Methylation.** Formation of *S*-methyl captopril was linear with respect to time when incubated with pET21 METTL7B. Simple linear regression analysis gives an R squared value of 0.99. All data is presented as the mean ± standard deviation.

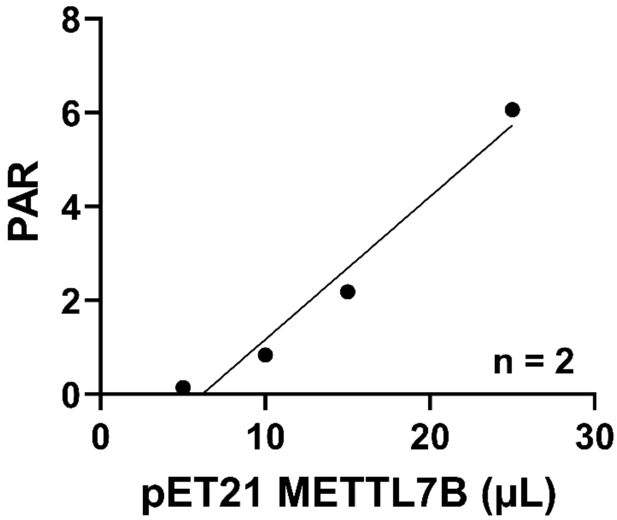


**Supplemental Data Figure 3: Protein Linearity of pET21 METTL7B Captopril Methylation.** Formation of *S*-methyl captopril was linear with respect to protein concentration when incubated with pET21 METTL7B. Simple linear regression analysis gives an R squared value of 0.96. All data is presented as the mean ± standard deviation.

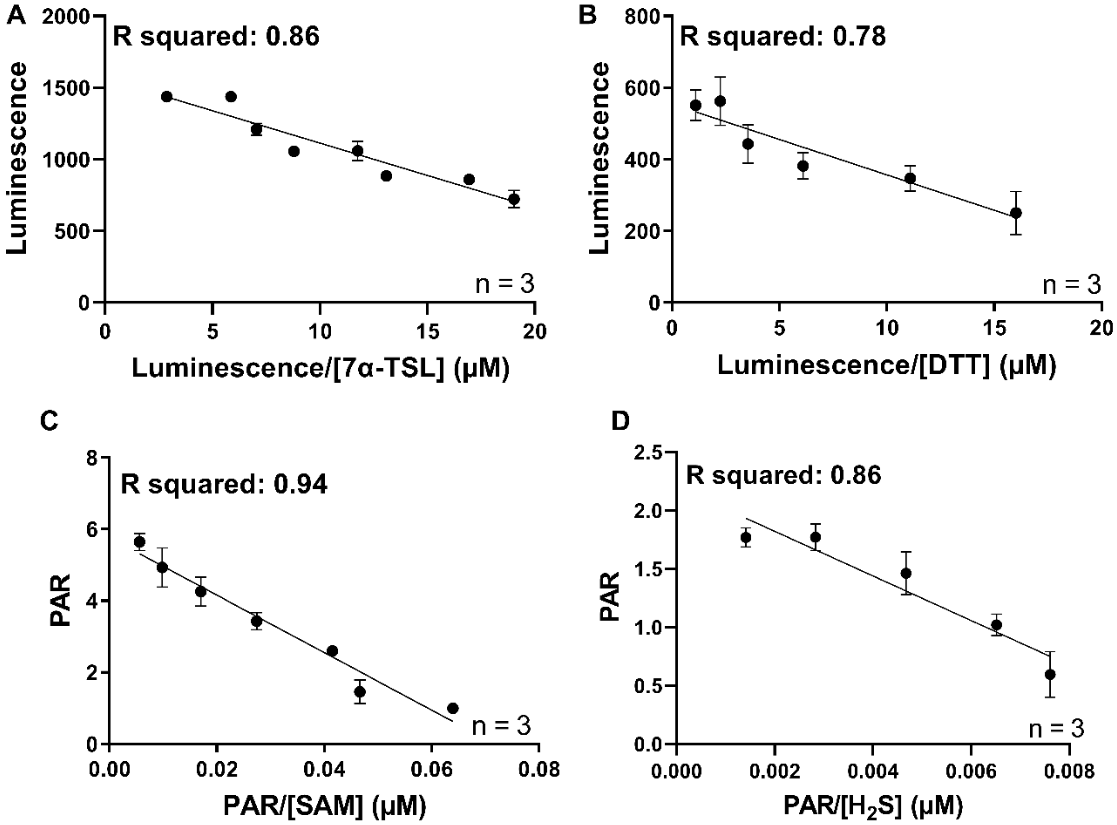


**Supplemental Data Figure 4: Eadie-Hofstee Transformations of thiol methylation by pET21 METTL7B. A)** Linearization of 7α-thiospironolactone methylation as measured by SAH formation. **B)** Linearization of dithiothreitol methylation as measured by SAH formation. **C)** Linearization of S-adenosyl-L-methionine use as measured by captopril methylation. **D)** Linearization of hydrogen sulfide methylation as measured by formation of methanethiol. All data is presented as the mean ± standard deviation.
